# Radiological Patterns in Maxillofacial Ewing Sarcoma: A systematic review with pooled patient-level descriptive analysis

**DOI:** 10.64898/2026.08.12.744464

**Authors:** Ajo Babu George, Shishir Maharana, Ridhi Agarwal, Abiya Mariam George, Sonam Khurana

## Abstract

**Background:** Ewing sarcoma (ES) is a rare malignant bone tumor with predilection for the mandible and maxilla in the head and neck region. However, existing literature comprises fragmented case reports and small case series that fail to establish consolidated, evidence-based understanding of characteristic radiological patterns in the maxillofacial region, hindering timely diagnosis and potentially leading to misdiagnosis or delayed intervention.

**Methodology:** A systematic review and pooled patient-level descriptive analysis were conducted according to the PRISMA guidelines and pre-registered on PROSPERO. Comprehensive searches of PubMed, OVID, and Cochrane databases (inception to July 2025) identified studies reporting radiological findings of biopsy-confirmed maxillofacial Ewing sarcoma. Quality assessment using Joanna-Briggs Institute criteria ensured inclusion of only high-quality cases (quality score ≥4/5). Synthesis Without Meta-analysis (SWiM) methodology with pooled prevalence estimation and binomial vote-counting analysis were employed for 68 published cases.

**Results:** Four radiological features demonstrated consistent predominance across pooled cases: soft tissue mass presence (100%, 95% CI: 94.7–100.0%), enhancing soft tissue (77.6%, 95% CI: 65.8–86.9%), cortical destruction (69.0%, 95% CI: 55.5–80.5%), and notably, absence of periosteal reaction (84.7%, 95% CI: 73.0–92.8%). Location-specific radiological phenotypes were evident: maxillary tumors demonstrated near-universal sinus involvement (100%) with high soft tissue enhancement (92.3%), whereas mandibular tumors showed predominant cortical destruction (80.0%) and teeth involvement (81.2%).

**Conclusion:** MRI and CT are essential for characterizing the distinctive radiological profile of maxillofacial Ewing sarcoma, enabling early identification and improving patient outcomes in this rare malignancy.

**Highlights:**

- First pooled review to summarize imaging features of maxillofacial Ewing sarcoma
- Analysis of 68 published cases reveals consistent imaging patterns.
- Most tumors show soft tissue mass and bone damage without surface reaction.
- Jaw tumors differ from long bone tumors in their imaging appearance.
- Upper and lower jaw tumors show distinct location-specific features.

## 1. Introduction

Ewing sarcoma (ES) is a rare malignant small round blue cell tumor described by James Ewing in 1921. It can occur in any bone but commonly it is found in the diaphysis of long bones and pelvic girdle. In the head and neck region, although rare, predilection is toward mandible followed by maxilla.^[1]^

It accounts for 10-15% of all primary malignant bone tumors affecting adolescents and young adults and it seldom develops after 30 years of age.^[2]^ The mean age of occurrence in the head and neck region is 10.9 years. ES generally affects white population and the male sex (male/female ratio, 1.3-1.5:1).^[3]^

According to anatomical site of occurrence it is classified as:(a) intraosseous (most common) (b) extraskeletal (less common) and (c) periosteal (rare).^[4]^

The majority of the available literature on the subject comprises single clinical cases. ^[5]^

The sporadic case reports, along with very small case series in many instances, however, fail to differentiate between the exact primary tumor locations by classifying “head and neck” as a collective potpourri, ^[6]^ leading to a lack of consolidated, evidence-based understanding of its characteristic radiological patterns. This fragmented knowledge hinders timely diagnosis and can lead to misdiagnosis or delayed intervention, with potentially severe consequences for patients. There is a critical gap in the existing oncology and radiology literature for a systematic synthesis and quantitative analysis of these radiological features.

This systematic review with pooled patient-level descriptive analysis aims to address this critical knowledge gap by systematically identifying, describing, and quantifying the characteristic radiological patterns of Ewing sarcoma in the maxillofacial region. By pooling data from published literature, we seek to establish more robust diagnostic criteria specific to this anatomical site, ultimately aiding radiologists and clinicians in early identification and improving patient outcomes.

## 2. Materials and Methods

### 2.1. Search Strategy and Selection Criteria

The systematic review was pre-registered on PROSPERO and was carried out following a predefined protocol that was used to perform this systematic review and descriptive analysis in accordance with the preferred reporting items for systematic reviews and meta-analyses (PRISMA) checklist. ^[7]^ The PubMed, OVFT, and Cochrane databases were searched to identify studies reporting radiological patterns of maxillofacial Ewing sarcoma from their inception to July 2025. We used the following search terms: “Ewing sarcoma”, “Ewing’s sarcoma”, “maxillofacial”, “jaw”, “mandible”, “maxilla” in conjunction with “radiology”, “imaging”, “computed tomography”, “magnetic resonance imaging”, “X-ray”, “positron emission tomography”, “bone destruction”, “periosteal reaction”, and “soft tissue mass”.

Successive use of Boolean operators [AND, OR] was also employed. The references of all the studies were screened to include relevant additional publications.

For descriptive analysis on the prevalence of radiological patterns in maxillofacial Ewing sarcoma, the inclusion criteria were:

1. Original case reports, case series, retrospective studies, and prospective studies; 2. Studies reporting imaging findings (conventional X-ray, CT, MRI, PET) of biopsy-confirmed Ewing sarcoma involving maxillofacial skeleton (maxilla, mandible, zygoma, nasal bones, paranasal sinuses, and other facial bones); 3. Studies published in English.

The exclusion criteria involved: 1. Review articles, editorials, conference abstracts without sufficient primary data, and book chapters; 2. Studies not providing clear descriptions or images of radiological features; 3. Studies involving non-maxillofacial Ewing sarcoma; 4. Non-English language articles where translation was not feasible.

## 3. Data Extraction and Analysis

### 3.1. Methods

A standardized, pre-piloted data extraction form was used to meticulously extract relevant data from each included study. Information extracted included Patient demographics: Age, sex; Tumor characteristics: Specific maxillofacial site, size; Imaging modality: Type of imaging used (X-ray, CT, MRI); Radiological features: Detailed description of patterns, including but not limited to: Osteolytic patterns (e.g., permeative, moth-eaten, geographic), Periosteal reactions (e.g., onion-skin, sunburst, spiculated, lamellated, Codman’s triangle), Cortical involvement (e.g., expansion, erosion, destruction, breach), Presence and characteristics of soft-tissue mass, Involvement of adjacent structures (e.g., teeth, nerves, sinuses) and Presence of sclerosis or calcifications.

In accordance with the Synthesis Without Meta-analysis (SWiM) guideline ^[8]^, we employed pooled prevalence estimation and vote counting based on direction of effect as synthesis methods, as conventional meta-analysis was not feasible due to the nature of the evidence (individual case reports and sparse case series without comparative data).

For each radiological feature, prevalence was calculated as the number of cases with the feature present divided by the total number of cases with available data for that feature, expressed as a percentage. Exact binomial confidence intervals (Clopper-Pearson method) ^[8]^ were calculated for all prevalence estimates, providing conservative estimates appropriate for case series data with modest to moderate sample sizes.

Vote counting was conducted to assess whether each radiological feature was present significantly more (or less) often than would occur by random chance. We applied one-tailed binomial tests for each feature, testing the null hypothesis that the feature would be present in 50% of cases by chance alone. Features with P-values <0.05 were considered to show statistically significant evidence of characteristic occurrence or absence. The 50% prevalence threshold used for binomial vote-counting was selected as a neutral reference point to assess feature predominance rather than as a biologically meaningful cutoff. This approach was used to support descriptive interpretation of pooled case-level data and does not imply population-level inference.

All radiological features were organized into six categories (osteolytic patterns, periosteal reactions, cortical changes, soft tissue characteristics, adjacent structure involvement, and other features). Features with insufficient data were excluded from denominators for the specific features involved, with sample sizes reported transparently for each analysis. To investigate potential heterogeneity in radiological presentation, we conducted stratified analyses comparing features across age groups (pediatric vs. adult) and primary anatomical sites. Pediatric cases were defined as patients aged ≤18 years, while adult cases were defined as >18 years.

Statistical analyses were performed using R (version 4.5.1) with packages tidyverse (version 2.0.0), ggplot2 (version 4.0.0), binom, reshape2, and gridExtra.

### 3.2. Quality Assessment

Quality was assessed using the Joanna-Briggs Institute (JBI) tools for case reports/series ^[10]^ by two independent reviewers (S.M. and R.A.) and disagreements were resolved by consultation with a third reviewer (A.B.G.).

Four domains were specially emphasized upon while assessing the quality of data: (1) adequacy of case definition (selection), (2) clarity of diagnostic establishment (ascertainment), (3) completeness of radiological feature reporting, and (4) adequacy of imaging quality. Each domain was rated as low risk, some concerns, or high risk based on predefined quality thresholds. Only cases rated as low risk across all domains (quality score ≥4) were included in the final analysis, ensuring methodological rigor.

Risk of bias assessment demonstrated low risk across all four evaluated domains for all 68 included cases (100%), reflecting the stringent inclusion criteria applied (minimum quality score 4/5). This indicates high confidence in the reliability of extracted radiological data and minimizes potential selection and reporting biases.

## 4. Statistics

### 4.1. Results

Among 68 cases of maxillofacial Ewing sarcoma with available radiological data, soft tissue mass was universally present (100%, 95% CI: 94.7-100.0%, n=68). This finding likely reflects both the true aggressive biological behavior of maxillofacial Ewing sarcoma and selective reporting bias, as cases lacking an appreciable soft tissue component may be less likely to be reported or imaged with advanced cross-sectional modalities. Cortical destruction was the most common osseous finding, present in 40 of 58 cases with reported data (69.0%, 95% CI: 55.5-80.5%, with statistically significant evidence for its characteristic occurrence when tested against the null hypothesis of 50% prevalence (binomial test, P=0.0054). The studies’ characteristics are shown in **Table 1**.

**Table 1.**
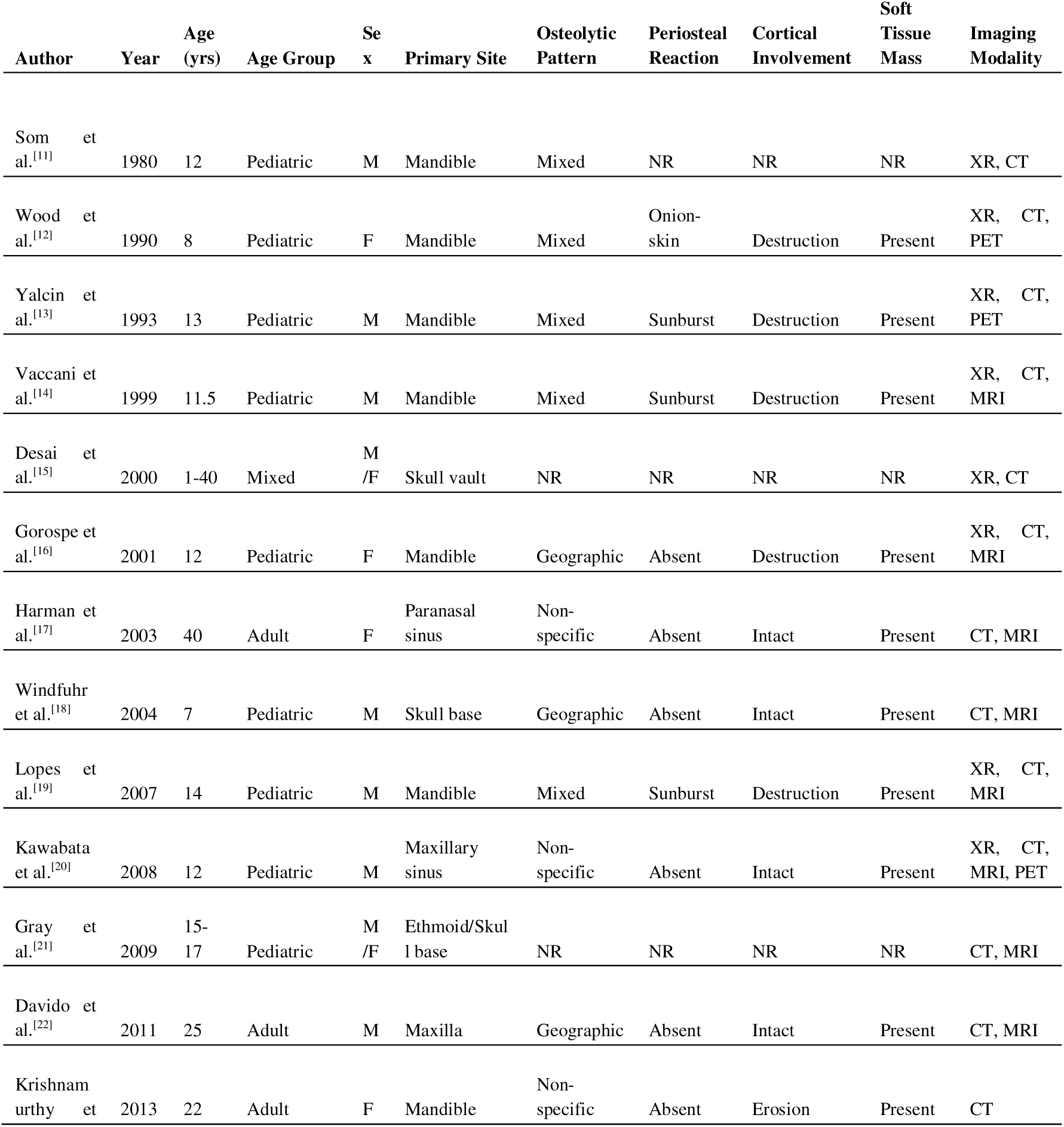
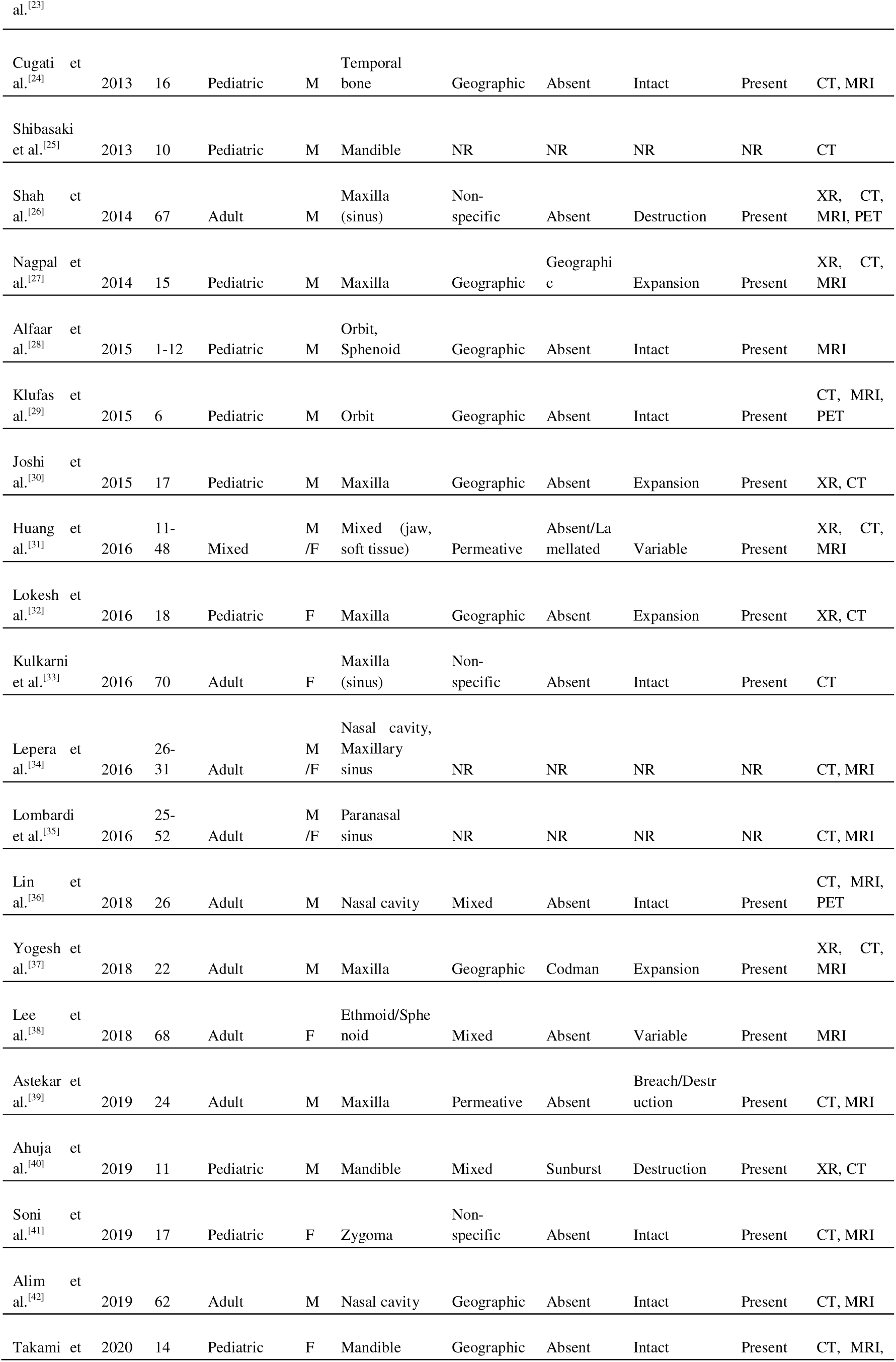
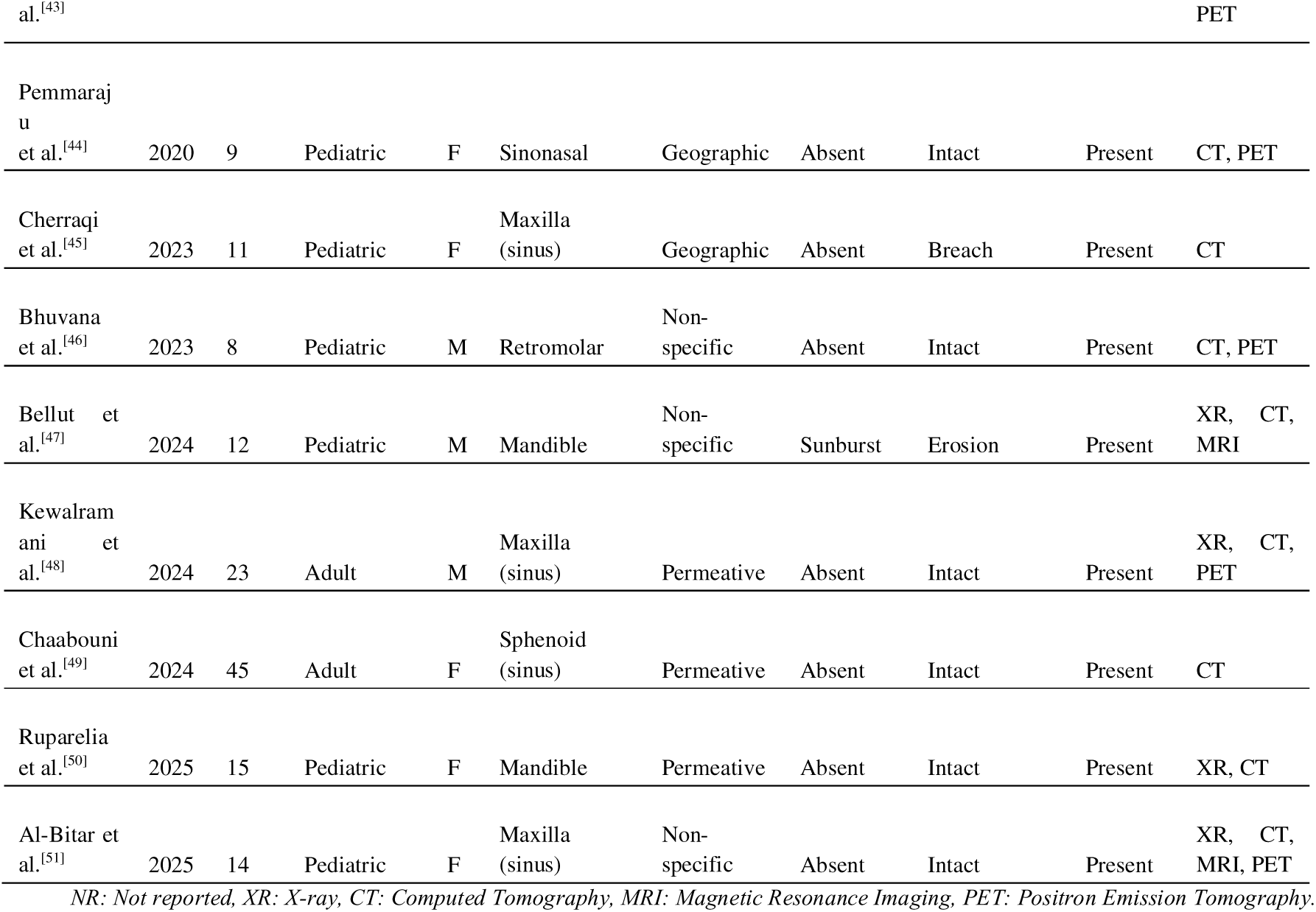
Literature Review of Maxillofacial Ewing Sarcoma–Clinical & Radiological Characteristics (41 Studies, 68 Cases, 1980-2025)

Periosteal reaction patterns showed the reverse phenomenon: the absence of periosteal reaction was the typical pattern, observed in 50 of 59 cases (84.7%, 95% CI: 73.0-92.8%, P<0.0001). When periosteal reaction was identified (9 cases), the sunburst pattern was most common (7/59, 11.9%, 95% CI: 4.9-22.9%), while other periosteal patterns (onionskin, lamellated, codman, spiculated) were rare (<5% each).

Soft tissue characteristics were highly consistent: not only was soft tissue mass universal, but enhancing soft tissue was present in 52 of 67 cases with available data (77.6%, 95% CI: 65.8-86.9%, P<0.0001), establishing this as a characteristic finding. Well-defined soft tissue margins were rare (2.9%, 95% CI: 0.4-10.2%).

Osteolytic patterns showed variable prevalence, with geographic pattern being most common (16/58, 27.6%, 95% CI: 16.7-40.9%), followed by permeative pattern (12/58, 20.7%, 95% CI: 11.2-33.4%). The moth-eaten pattern and cases with no osteolytic changes were both uncommon (5.2% each).

Adjacent structure involvement varied moderately: sinus involvement (44.1%, n=30/68), teeth involvement (32.4%, n=22/68), muscle involvement (17.6%, n=12/68), orbital involvement (16.2%, n=11/68), and nerve involvement (8.8%, n=6/68). Pathologic fracture was not observed in any case (0%, 95% CI: 0-5.3%). A comprehensive list of the features examined is revealed in **Table 2**.

**Table 2.**
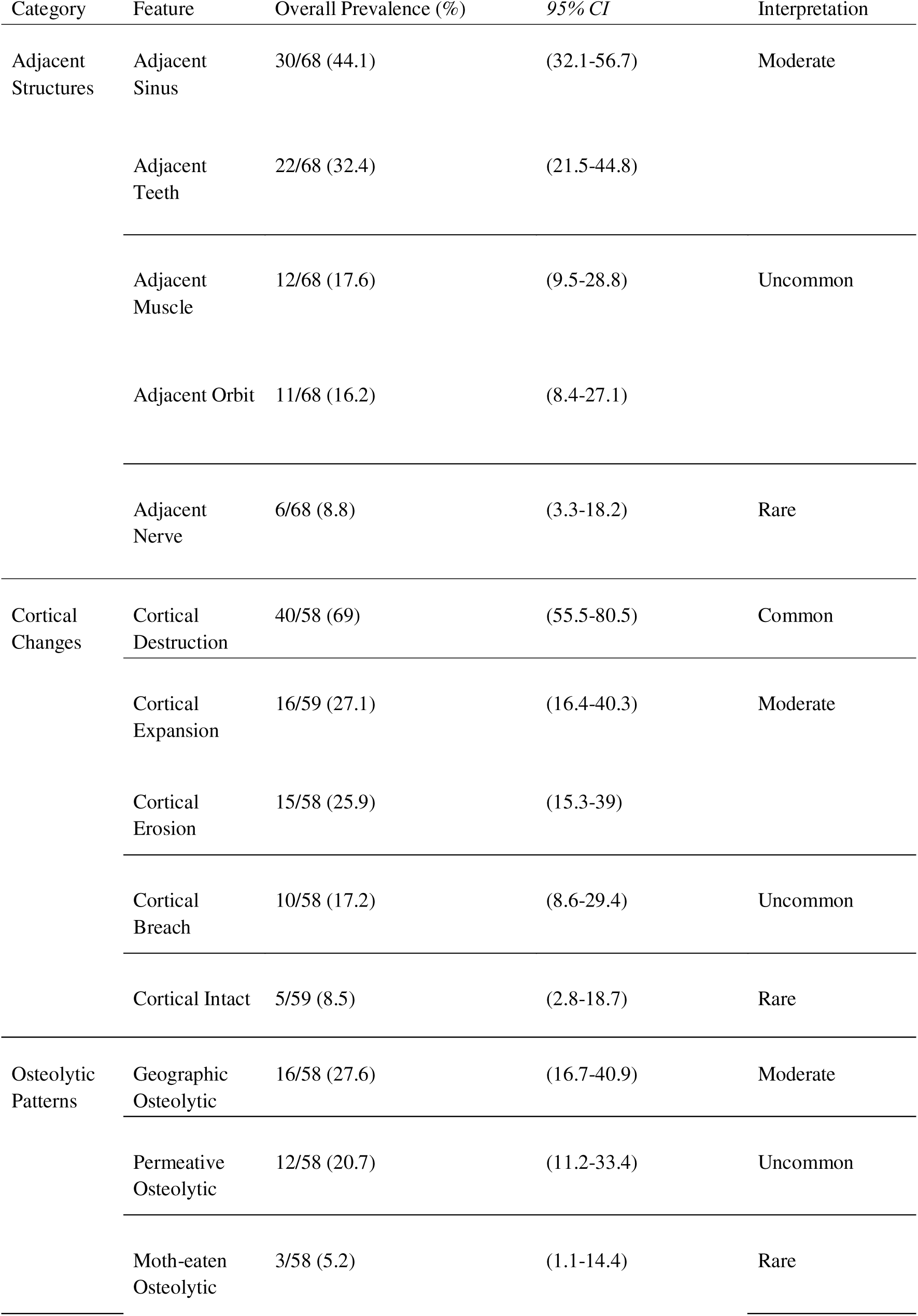
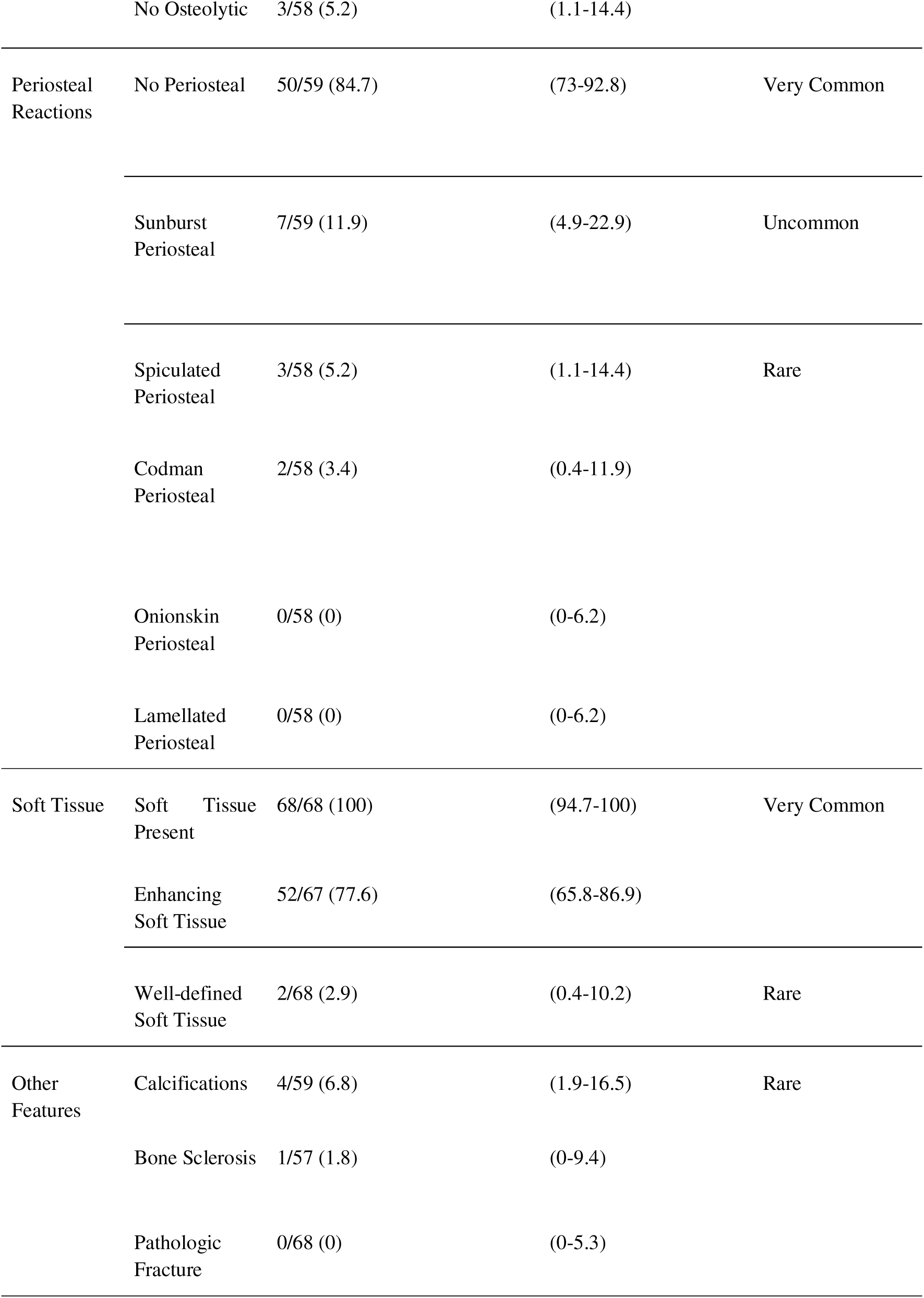
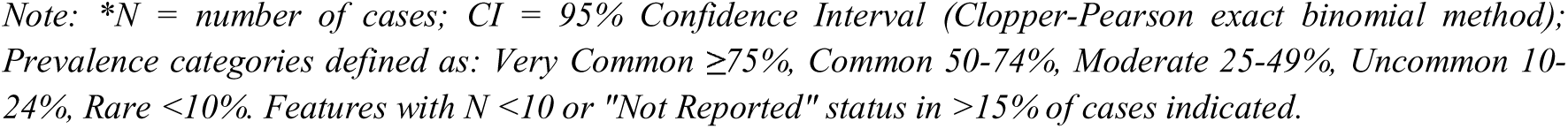
Prevalence of Various Radiological Features of Ewing Sarcoma.

| Category | Feature | Overall Prevalence (%) | 95% *CI* | Interpretation |
| --- | --- | --- | --- | --- |
| Adjacent Structures | Adjacent Sinus | 30/68 (44.1) | (32.1-56.7) | Moderate |
|  | Adjacent Teeth | 22/68 (32.4) | (21.5-44.8) |  |
|  | Adjacent Muscle | 12/68 (17.6) | (9.5-28.8) | Uncommon |
|  | Adjacent Orbit | 11/68 (16.2) | (8.4-27.1) |  |
|  | Adjacent Nerve | 6/68 (8.8) | (3.3-18.2) | Rare |
| Cortical Changes | Cortical Destruction | 40/58 (69) | (55.5-80.5) | Common |
|  | Cortical Expansion | 16/59 (27.1) | (16.4-40.3) | Moderate |
|  | Cortical Erosion | 15/58 (25.9) | (15.3-39) |  |
|  | Cortical Breach | 10/58 (17.2) | (8.6-29.4) | Uncommon |
|  | Cortical Intact | 5/59 (8.5) | (2.8-18.7) | Rare |
| Osteolytic Patterns | Geographic Osteolytic | 16/58 (27.6) | (16.7-40.9) | Moderate |
|  | Permeative Osteolytic | 12/58 (20.7) | (11.2-33.4) | Uncommon |
|  | Moth-eaten Osteolytic | 3/58 (5.2) | (1.1-14.4) | Rare |
|  | No Osteolytic | 3/58 (5.2) | (1.1-14.4) |  |
| Periosteal Reactions | No Periosteal | 50/59 (84.7) | (73-92.8) | Very Common |
|  | Sunburst Periosteal | 7/59 (11.9) | (4.9-22.9) | Uncommon |
|  | Spiculated Periosteal | 3/58 (5.2) | (1.1-14.4) | Rare |
|  | Codman Periosteal | 2/58 (3.4) | (0.4-11.9) |  |
|  | Onionskin Periosteal | 0/58 (0) | (0-6.2) |  |
|  | Lamellated Periosteal | 0/58 (0) | (0-6.2) |  |
| Soft Tissue | Soft Tissue Present | 68/68 (100) | (94.7-100) | Very Common |
|  | Enhancing Soft Tissue | 52/67 (77.6) | (65.8-86.9) |  |
|  | Well-defined Soft Tissue | 2/68 (2.9) | (0.4-10.2) | Rare |
| Other Features | Calcifications | 4/59 (6.8) | (1.9-16.5) | Rare |
|  | Bone Sclerosis | 1/57 (1.8) | (0-9.4) |  |
|  | Pathologic Fracture | 0/68 (0) | (0-5.3) |  |

Note: \*N = number of cases; CI = 95% Confidence Interval (Clopper-Pearson exact binomial method); Prevalence categories defined as: Very Common $\geq 75\%$ , Common 50-74%, Moderate 25-49%, Uncommon 10-24%, Rare $<10\%$ . Features with N <10 or "Not Reported" status in >15% of cases indicated.

### 4.2. Distribution

A visual summary of the presence/absence and reporting completeness distribution of 10 key radiological features across 68 cases of maxillofacial Ewing sarcoma is depicted in **Figure 2**. Soft tissue findings were universally present, while cortical and osteolytic patterns showed heterogeneous color distribution (varying prevalence).

**Figure 1.**
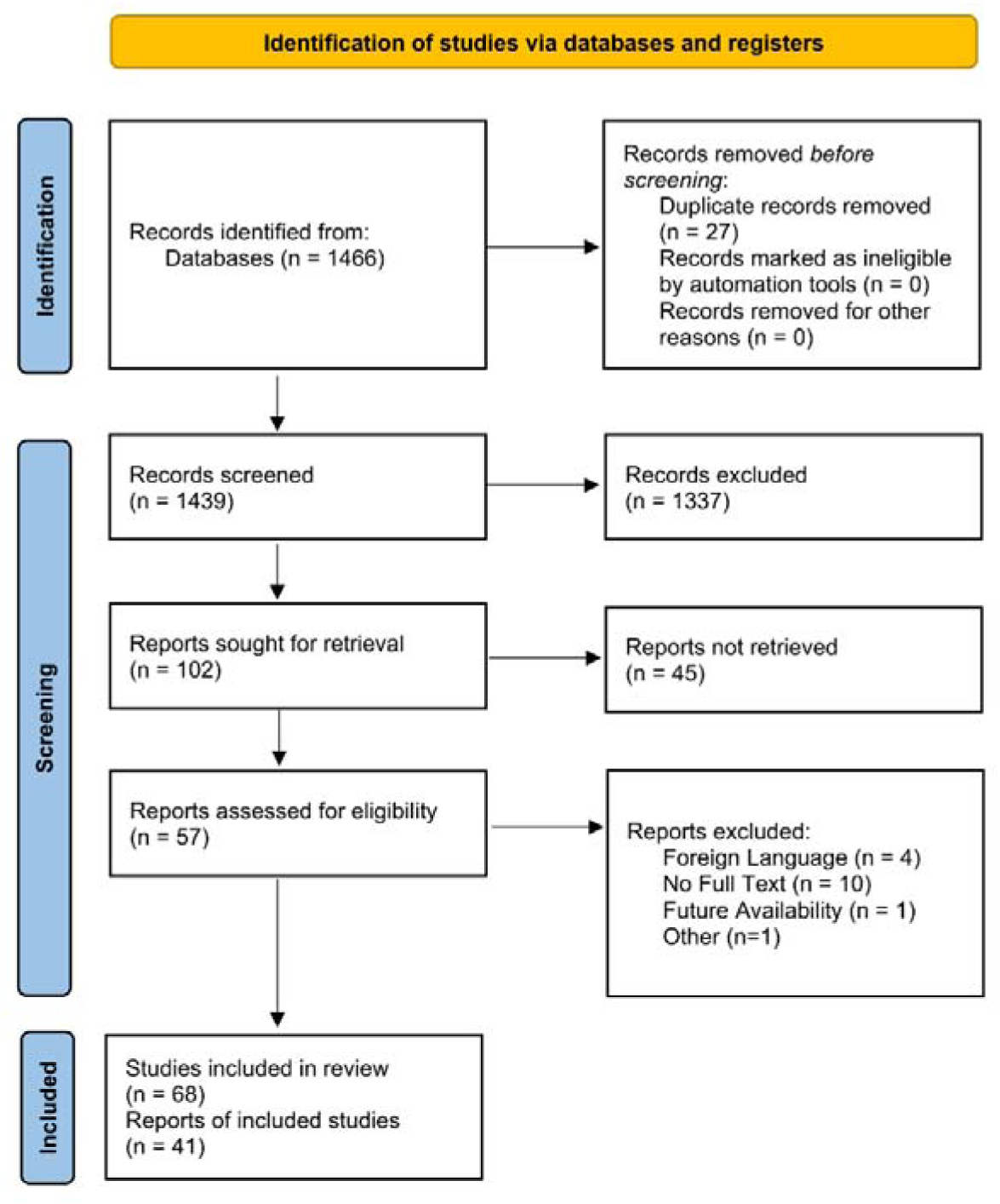
The PRISMA flow diagram.

**Figure 2.**
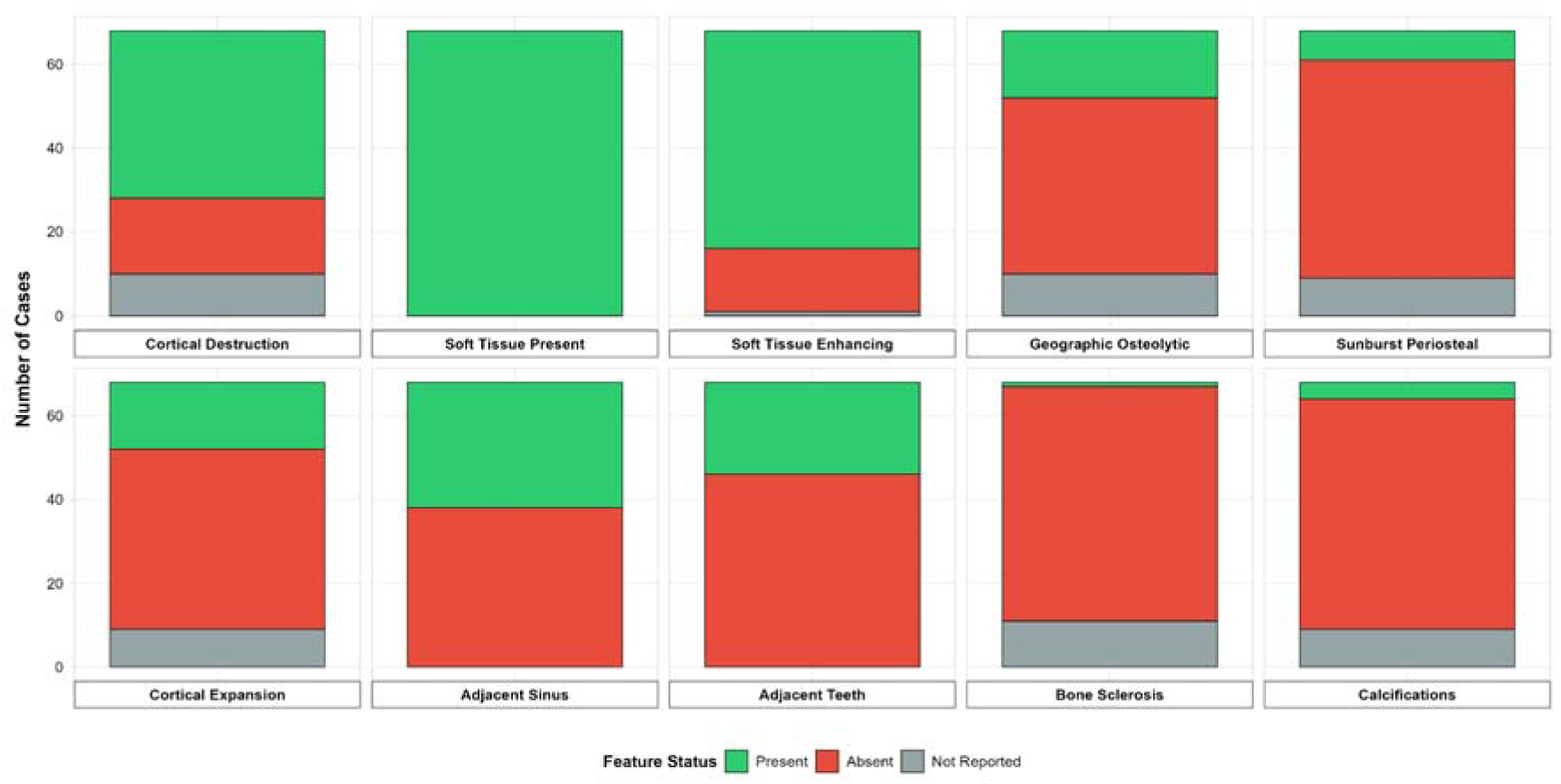
A harvest plot summarising prevalence of key features. The stacked bar height indicates total case count, while the color composition demonstrates both feature prevalence and reporting completeness.

### 4.3. Subgroup Analysis

**Figure 3(a)** reveals modest variation between pediatric and adult cases. Cortical destruction occurred at similar frequencies in both age groups (pediatric 68.4% vs. adult 70.0%).

**Figure 3.**
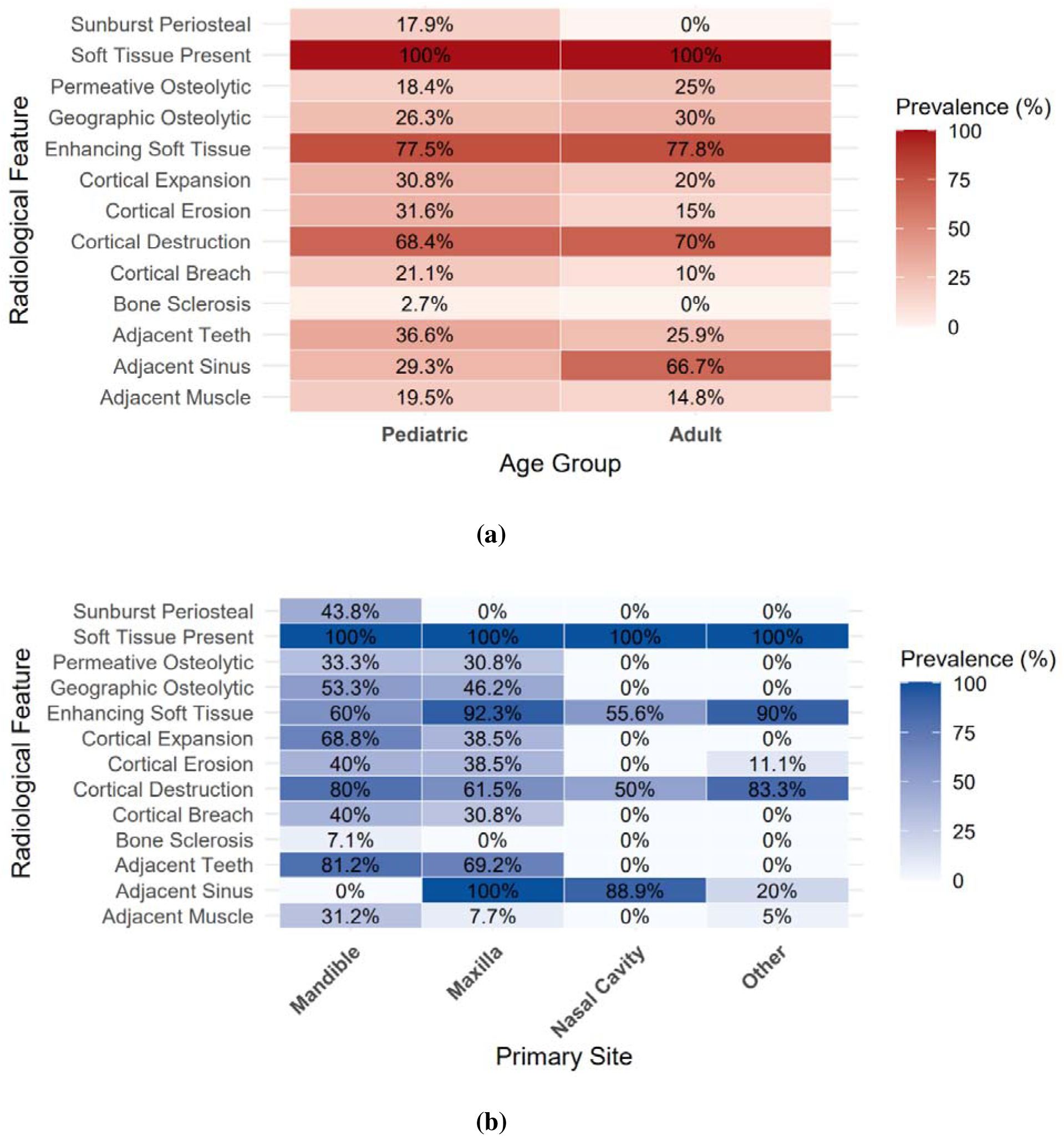
**(a)** Heat map comparing prevalence of radiological features between pediatric cases (n=41, red gradient) and adult cases (n=27). **(b)** Heat map comparing feature prevalence across primary anatomical sites (mandible n=16, maxilla n=13, nasal cavity n=9, other n=20).

However, cortical erosion was more common in pediatric cases (31.6% vs. 15.0%), as was periosteal sunburst pattern (17.9% vs. 0.0%, exclusively pediatric). Conversely, sinus involvement was substantially more common in adult cases (66.7% vs. 29.3%), with the difference being clinically notable (37.4 percentage point difference).

**Figure 3(b)** casts light on the marked influence of primary tumor location on radiological presentation patterns. Maxillary tumors (n=13) demonstrated near-universal sinus involvement (100%) and high enhancing soft tissue prevalence (92.3%), consistent with the proximity of the tumor to the maxillary sinus. Mandibular tumors (n=16) showed predominant cortical involvement (cortical destruction 80.0%, cortical expansion 68.8%) and notably high teeth involvement (81.2%), reflecting the anatomical relationship between mandibular tumors and tooth roots. Nasal cavity tumors (n=9) similarly showed high sinus involvement (88.9%), whereas mandibular cases showed no sinus involvement (0%). Other sites (n=20) showed the highest cortical destruction rate (83.3%) but low sinus involvement (20.0%).

These patterns suggest distinct radiological phenotypes based on tumor location, which may have diagnostic implications for radiologists evaluating suspected maxillofacial Ewing sarcoma.

Notable modality-dependent differences are revealed in **Figure 4(a)**: soft tissue enhancement detected most frequently on CT (82.3%) and X-ray (82.1%) compared to MRI (74.3%) and PET (69.2%); cortical destruction prevalence ranges from 76.9% on X-ray to 54.5% on PET; geographic osteolytic patterns vary markedly (40.0% on MRI vs. 9.1% on PET); and soft tissue presence remains universal (100%) across all imaging modalities. These patterns reflect differential detection capabilities of imaging modalities, particularly soft tissue characterization. CT and MRI demonstrate superior soft tissue assessment, while osseous features show variable modality-dependent detection rates. Observed differences across imaging modalities likely reflect variation in detection capabilities rather than true biological differences, as many cases underwent multiple imaging techniques and absence of a feature on a given modality does not equate to true absence.

**Figure 4.**
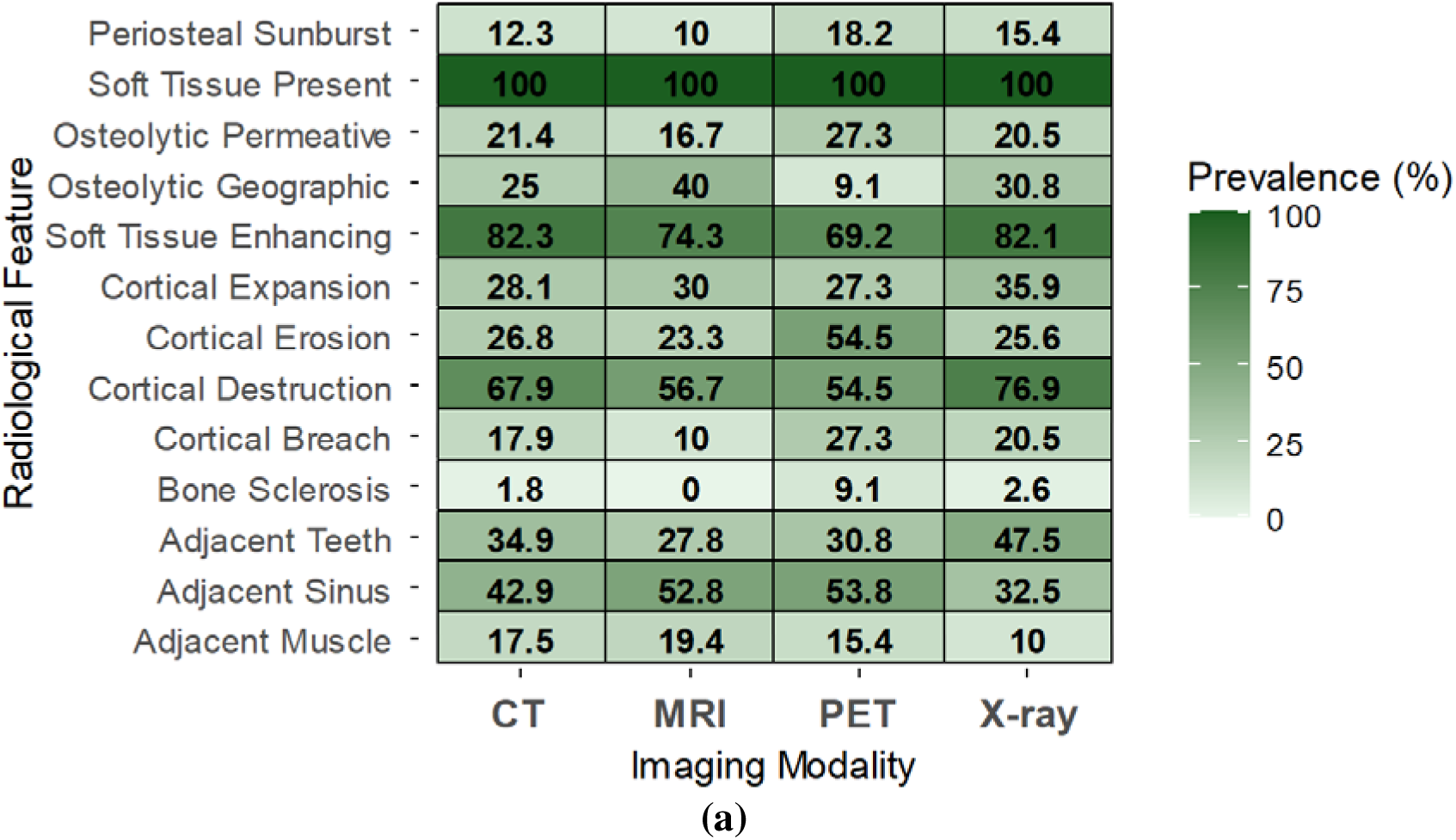
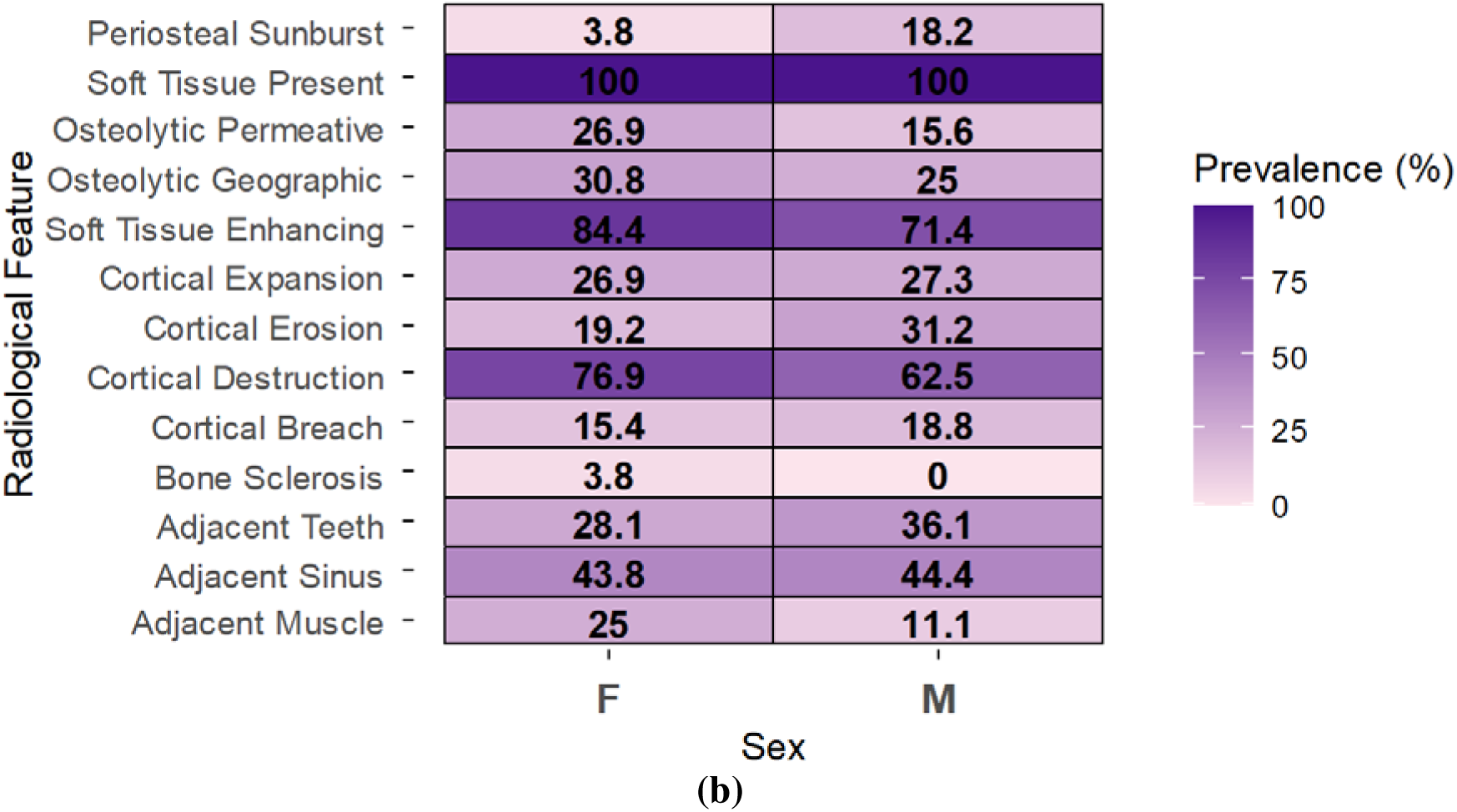
**(a)** Heat map comparing prevalence of radiological features across imaging modalities (CT n=63, MRI n=36, PET n=13, X-ray n=40). **(b)** Heat map comparing prevalence of radiological features between male cases (n=36, left column) and female cases (n=32, right column).

**Figure 4(b)** unveils sex-related differences, with periosteal sunburst reaction predominantly identified in males (18.2% vs. 3.8%); periosteal absence more prevalent in females (96.2% vs. 75.8%, difference −20.4%); cortical erosion more common in males (31.2% vs. 19.2%); and soft tissue enhancement higher in females (84.4% vs. 71.4%). However, most features show comparable prevalence between sexes with overlapping confidence intervals, including soft tissue presence (100% in both), cortical destruction (62.5% males vs. 76.9% females), and adjacent muscle involvement (25.0% females vs. 11.1% males). Overall, radiological presentation is generally similar between sexes with periosteal reaction patterns showing the most notable variation.

## 5. Discussion

The present systematic review identified four radiological features that consistently predominated across pooled published cases of maxillofacial Ewing sarcoma: (1) universal soft tissue mass presence (100%), (2) enhancing quality of soft tissue (77.6%), (3) typical absence of periosteal reaction (84.7%), and (4) cortical destruction in approximately two-thirds of cases (69.0%). These four features comprise the core radiological profile of maxillofacial Ewing sarcoma and should prompt clinical consideration of this diagnosis. However, this apparent universality should be interpreted cautiously, as reliance on published case reports may preferentially capture tumors with conspicuous extraosseous components, while lesions with subtle or minimal soft tissue extension may be underrepresented.

This pattern contrasts with the classic radiological appearance of long-bone Ewing sarcoma, which frequently presents with aggressive periosteal reactions (sunburst or onionskin patterns). The atypical presentation in maxillofacial ES with predominantly osseous destruction rather than periosteal reaction likely contributes to diagnostic delays, as radiologists and clinicians may not consider Ewing sarcoma in the absence of classic periosteal reactions.

Maxillary tumors present with a distinctive phenotype dominated by sinus involvement and enhancing soft tissue, whereas mandibular tumors show predominantly cortical changes with teeth involvement. These site-specific variations/patterns are clinically significant and should be recognized by radiologists to improve early detection and guide appropriate clinical follow-up. Recognition of these site-specific imaging phenotypes may assist radiologists in refining differential diagnoses when evaluating aggressive maxillofacial lesions, particularly in pediatric and young adult populations.

Overall, maxillofacial Ewing sarcoma demonstrates a distinctive radiological phenotype characterized by aggressive cortical destruction and prominent soft tissue mass, yet a paradoxical absence of periosteal reaction. Awareness of this atypical presentation is critical, as reliance on classic long-bone imaging criteria may delay diagnosis in the maxillofacial region. Early use of cross-sectional imaging, particularly CT and MRI, is essential to recognize these patterns and facilitate timely multidisciplinary oncologic management.

### 5.1. Limitations

This synthesis is based entirely on case reports and small case series, which are inherently subject to publication bias toward unusual or diagnostically instructive presentations. The universal prevalence of soft tissue mass and cortical destruction in our series may overestimate their true clinical frequency if cases with subtle or atypical presentations are underreported. Additionally, heterogeneity in imaging modalities (radiography, CT, MRI, PET), technical parameters, and reporting completeness across studies may have influenced prevalence estimates. Some radiological features were not uniformly reported, requiring us to exclude 10-11 cases from analyses of specific features; this selective reporting may introduce bias.

The analysis represents descriptive synthesis of individual patient data and cannot estimate true population prevalence or establish causality. Nevertheless, the systematic identification and transparent reporting of radiological patterns in maxillofacial ES provides valuable insights for diagnostic practice.

Because the analysis is based exclusively on published case reports and small case series, radiological features that are subtle, atypical, or clinically unrecognized may be underrepresented, potentially inflating the apparent prevalence of aggressive imaging findings.

## 6. Conclusions

This systematic review provides the first pooled patient-level synthesis defining the characteristic radiological phenotype of maxillofacial Ewing sarcoma, addressing a critical evidence gap in oral and maxillofacial oncology.

The analysis identifies four radiological features that consistently predominate in published cases of maxillofacial Ewing sarcoma: (1) universal soft tissue mass (100%), (2) enhancing soft tissue (77.6%), (3) cortical destruction (69.0%), and (4) notably, the absence of periosteal reaction (84.7%). This constellation represents a distinctive radiological phenotype that contrasts markedly with Ewing sarcoma in long bones, where aggressive periosteal reactions (sunburst or onionskin patterns) are characteristic.

The demonstration of location-specific radiological phenotypes has significant diagnostic implications. Maxillary tumors present with near-universal sinus involvement (100%) and high soft tissue enhancement (92.3%), suggesting that Ewing sarcoma should be considered in the differential diagnosis of sinonasal masses with aggressive bone destruction in young patients. Mandibular tumors, by contrast, manifest predominantly through cortical destruction and teeth involvement (81.2%), reflecting their characteristic intraosseous growth pattern and intimate relationship with dental structures.

This maxillofacial profile is atypical and contrasts markedly with Ewing sarcoma in long bones, where aggressive periosteal reactions (like sunburst or onionskin patterns) are characteristic. The relative absence of periosteal reaction in the jaw likely contributes to diagnostic delays and should prompt heightened clinical vigilance.

While cortical destruction occurs with similar frequency across age groups, the identification of periosteal sunburst reaction exclusively in pediatric cases suggests that this finding, when present, may indicate Ewing sarcoma in younger patients and warrants urgent evaluation.

Clinicians should recognize this specific four-feature profile (soft tissue mass, enhancement, cortical destruction, and absence of periosteal reaction) as suggestive of maxillofacial Ewing sarcoma. Appropriate diagnostic workup must include advanced cross-sectional imaging (CT and MRI) to assess these patterns, enabling timely referral for multidisciplinary oncologic management.

## Author Contributions

Conceptualization—A.B.G. and S.M.; methodology—A.B.G. and S.M.; software—S.M.; validation—A.M.G. and R.A.; formal analysis—S.M.; investigation—S.M. and R.A.; resources—A.B.G.; data curation—R.A.; writing—original draft preparation—A.B.G., S.M. and A.M.G. Manuscript writing—S.M.; visualization— S.M.; supervision—A.B.G., S.K.; and S.M.—project administration. All authors have read and agreed to the published version of the manuscript.

